# Simultaneous electrophysiological recording and fiber photometry in freely behaving mice

**DOI:** 10.1101/807602

**Authors:** Amisha A Patel, Niall McAlinden, Keith Mathieson, Shuzo Sakata

## Abstract

*In vivo* electrophysiology is the gold standard technique used to investigate sub-second neural dynamics in freely behaving animals. However, monitoring cell-type-specific population activity is not a trivial task. Over the last decade, fiber photometry based on genetically encoded calcium indicators has been widely adopted as a versatile tool to monitor cell-type-specific population activity *in vivo*. However, this approach suffers from low temporal resolution. Here, we combine these two approaches to monitor both sub-second field potentials and cell-type-specific population activity in freely behaving mice. By developing an economical custom-made system, and constructing a hybrid implant of an electrode and a fiber optic cannula, we simultaneously monitor artifact-free pontine field potentials and calcium transients in cholinergic neurons across the sleep-wake cycle. We find that pontine cholinergic activity co-occurs with sub-second pontine waves, called P-waves, during rapid eye movement sleep. Given the simplicity of our approach, simultaneous electrophysiological recording and cell-type-specific imaging provides a novel and valuable tool for interrogating state-dependent neural circuit dynamics *in vivo*.

## Introduction

Intracranial electrophysiological recordings monitor neuronal activity at various spatial scales, from single cells to populations across brain regions, with high temporal resolution (Buzsaki, 2004;Buzsaki et al., 2012;Jun et al., 2017). However, one of limitations in this approach is identifying the source of the neural signal: because neuronal activity is typically monitored extracellularly in freely behaving condition, the identification/isolation of recorded neurons is challenging (Einevoll et al., 2012;Harris et al., 2016).

Genetically encoded indicators offer complementary advantages over *in vivo* electrophysiological approaches (Lin and Schnitzer, 2016;Deo and Lavis, 2018;Wang et al., 2019). Over the last two decades, genetically encoded calcium indicators (GECIs) have been widely used to interrogate not only neuronal ensemble dynamics, but also activity of non-neuronal cells, such as astrocytes *in vivo* (Nakai et al., 2001;Chen et al., 2013;Stobart et al., 2018;Dana et al., 2019;Inoue et al., 2019;Stringer et al., 2019). For example, GECIs enable to target specific cell-types and allow for the long-term monitoring of neuronal activity *in vivo*. However, because of the intrinsic nature of calcium signals, the low temporal resolution of GECIs are not ideal for monitoring sub-second neural dynamics. It is also challenging to monitor individual neuronal activity in deep brain areas without causing significant tissue damage.

Here, we combine an electrophysiological approach with GECI-based fiber photometry to simultaneously monitor both field potentials and calcium transients in freely behaving mice. Fiber photometry is an imaging method used to monitor fluorescent signals via an implanted fiber optic cannula (Adelsberger et al., 2005;Lutcke et al., 2010;Kim et al., 2016;Sych et al., 2019). Although it is still invasive, the diameter of the cannula is thinner than an endoscope. Therefore, fiber photometry is well-suited to monitor neural population activity in deep tissue, such as the brainstem.

In the present study, we develop a versatile custom-made fiber photometry system with the capability to integrate *in vivo* electrophysiological recording for freely behaving mice. To validate the versatility of our system, we ask whether sub-second neural activity can be observed together with cell-type specific calcium transients in freely behaving mice. To this end, we note the fact that pontine (P) or ponto-geniculo-occipital (PGO) waves appear during rapid eye movement (REM) sleep as sub-second brain waves, and that pontine cholinergic neurons play a critical role in the generation of P-waves (Callaway et al., 1987;Datta, 1997;Tsunematsu et al., 2019). Therefore, we monitor calcium transients from GCaMP6s-expressing pontine cholinergic neurons across the sleep-wake cycle along with cortical electroencephalograms (cEEGs), electromyograms (EMGs), and pontine EEGs (pEEGs). We show that P-waves during REM sleep co-occurs with calcium transients in cholinergic neurons. Thus, our system allows simultaneous electrophysiological recording and fiber photometry in freely behaving mice.

## Materials and Methods

### Recording system configuration

The recording system is shown in **Figure 1** and a parts list for the fiber photometry system is summarized in **Table 1**. The fiber photometry system consisted of two excitation channels. A 470 nm LED (M470L3, Thorlabs) was used to extract a Ca^2+^-dependent signal and a 405 nm LED (M405L3, Thorlabs) was used to obtain a Ca^2+^- independent isosbestic signal. Light from the LEDs was directed through excitation filters (FB470-10, FB405-10, Thorlabs) and a dichroic mirror to the fiber launch (DMLP425R and KT110/M, respectively). The fiber launch was connected to a multimode patch cable (M82L01, Thorlabs) which could be reversibly attached and detached to an implantable optic fiber on the mouse via a ceramic mating sleeve (CFM14L05 and ADAF1, respectively). Light emissions from GCaMP6s expressing neurons were then collected back through the optic fiber, and directed through a detection path, passing a dichroic mirror (MD498) to reach a photodetector (NewFocus 2151, Newport). A National Instruments DAQ (NI USB-6211) and custom-written LabVIEW software was used to control the LEDs and acquire fluorescence data at 1 kHz. LEDs were alternately turned on and off at 40 Hz in a square pulse pattern. Electrophysiology signals were recorded at 1 kHz using an interface board (RHD2000, Intan Technologies) and connected to the mouse via an amplifier (RHD2132 16-channel amplifier board, Intan Technologies).

**Table 1.** Parts list for the fiber photometry system.

| Component | Supplier | Product Code | Quantity |
| --- | --- | --- | --- |
| Photodetector | Newport | NewFocus 2151 | 1 |
| 470nm LED | Thorlabs | M470L3 | 1 |
| 405nm LED |  | M405L3 | 1 |
| LED driver |  | LEDD1B | 2 |
| LED power source |  | KSP101 | 2 |
| LED holder |  | CP12 | 2 |
| Dichroic mirror |  | DMLP425R | 1 |
| Dichroic mirror |  | MD498 | 1 |
| Excitation filter |  | FB470-10 | 1 |
| Excitation filter |  | FB405-10 | 1 |
| Bandpass filter |  | MF525-39 | 1 |
| Broadband dielectric mirror |  | BB1-E02 | 2 |
| Mirror mount |  | KCB1C/M | 2 |
| Aspheric lens |  | AL2520M-A | 4 |
| Cage plate for lens |  | CP08/M | 4 |
| Lens tubes |  | SM1L03-P5 | 5pk |
| Filter holder |  | C4W | 2 |
| Filter holder |  | B4C/M | 2 |
| Filter holder |  | FFM1 | 2 |
| Fiber launch |  | KT110/M | 1 |
| Patch cable |  | M82L01 | 1 |
| Optic fiber implant |  | CFM14L05-10 | 10pk |
| Mating sleeve |  | ADAF1-5 | 5pk |

**Figure 1.**
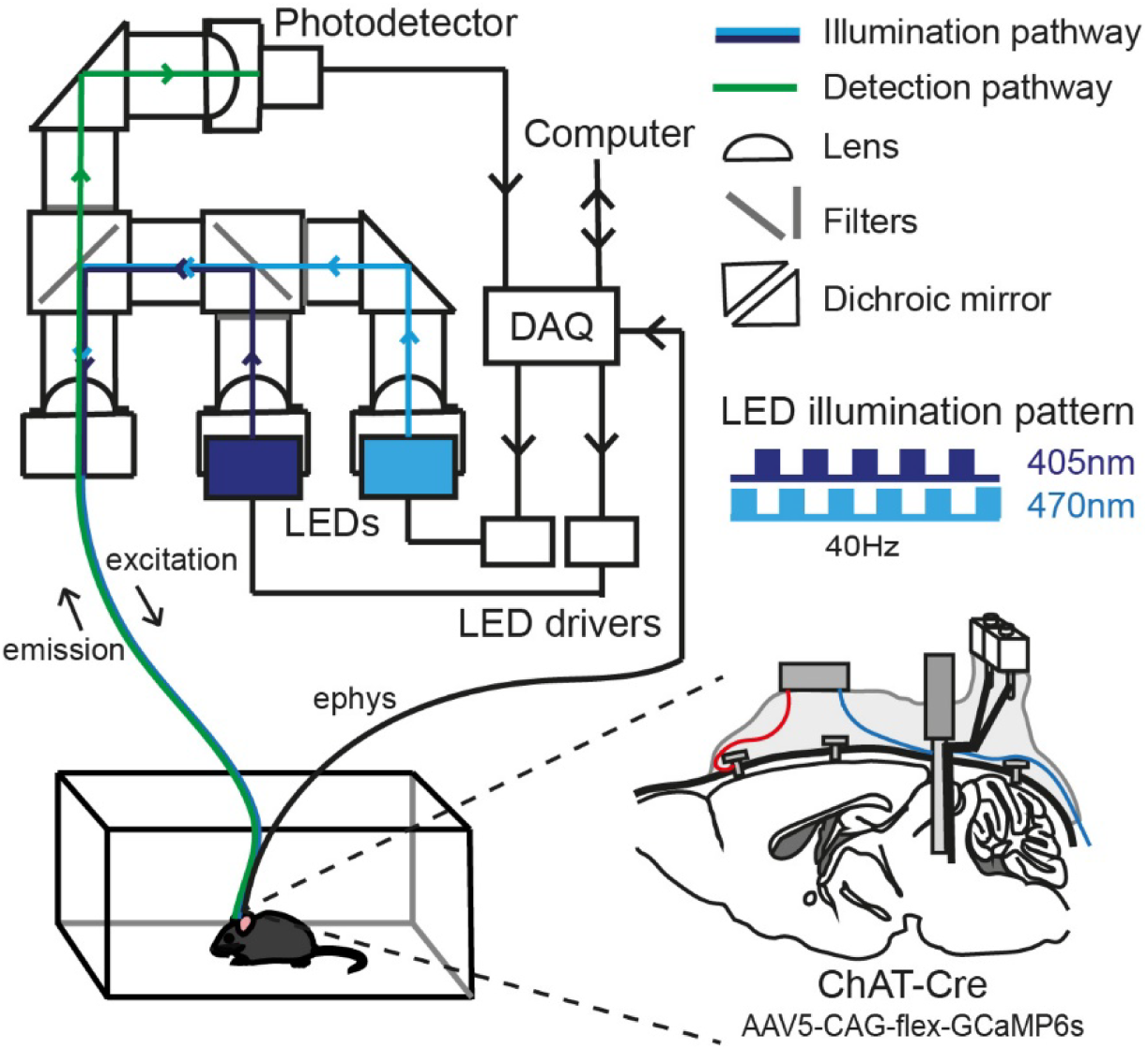
An integrated system of fiber photometry and electrophysiology. A schematic drawing (*top left*) provides the design of the fiber photometry system with two LEDs. The detailed parts list is provided in **Table 1**. The data acquisition (DAQ) module provides analogue outputs to drive the LEDs with a 40 Hz alternated illumination pattern, and obtains analogue inputs from the photodetector for fluorescent signals and electrophysiological signals, which are processed on an Intan Technologies’ system (not shown). Another schematic drawing (*bottom right*) illustrates the implant configuration: cortical EEGs were monitored via bone screws implanted over the frontal cortex (red). EMGs were monitored via twisted wires (blue) inserted to the neck muscle. The implant is a hybrid bipolar electrode and optical fiber. In this study, we expressed GCaMP6s in pontine ChAT+ neurons.

### Implant fabrication

Hybrid implants (also referred to as an optrode) consisted of a bipolar electrode (pair of electrodes) glued to the optic fiber and were fabricated though a multistep process. First two 0.1 mm diameter stainless steel wires (FE205850/2, Goodfellow) were glued together (offset by approximately 0.5 - 1 mm at the tip). Insulation was removed at one end and connected to a 2-piece connector (SS-132-T-2-N, Samtec) using conductive epoxy (186-3593, RS-Pro). The conductive epoxy was left to dry for 10 minutes and then secured with dental cement. Impedances were checked by connecting the bipolar electrodes to the Intan system (RHD2132 16-channel amplifier board and RHD2000, Intan Technologies) with a custom-made connector, and placing the tips of the bipolar electrodes in saline. Electrodes with impedances between 200 kΩ and 1 MΩ at 1 kHz were folded (as shown in **Fig. 2A**) and positioned alongside the optic fiber. The electrodes were fixed in place 500 µm below the tip of the optic fiber with superglue (473-455, RS-Pro), taking care not to get glue on the tips of either the optic fiber or bipolar electrode. Dental cement was then used to secure and stabilize the structure. Impedances were checked again (the range was 276 – 452 kΩ) and optrodes were ready for implantation.

**Figure 2.**
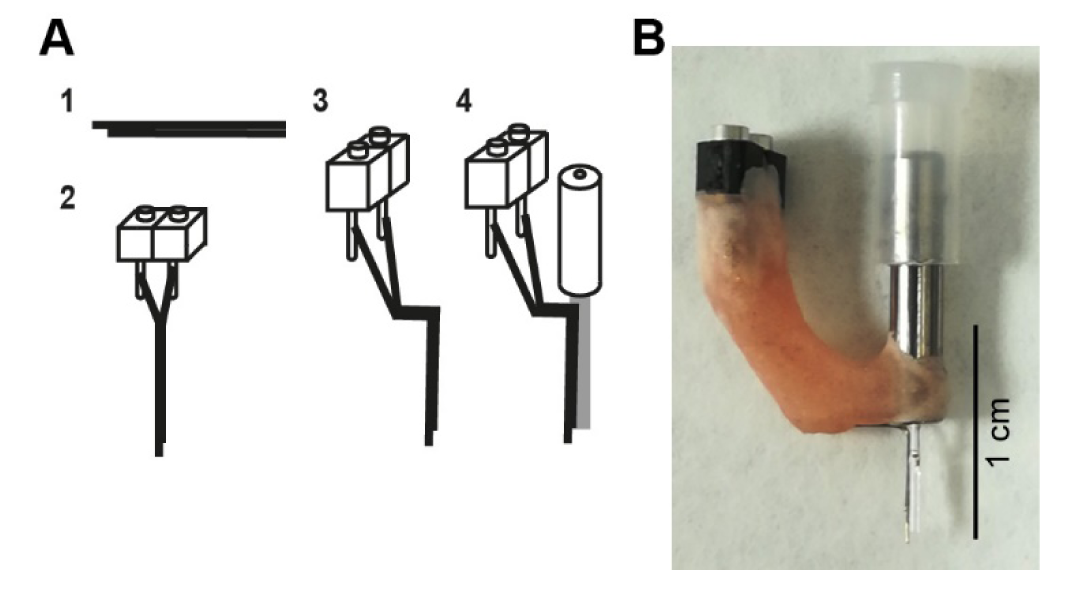
Hybrid implant. (**A**)The fabrication process for making hybrid implants. (1) Two wires are glued together to create a bipolar electrode. To differentiate local signals, the tip of two wires was separated by up to 1 mm. (2) The bipolar electrode is attached to a connector. (3) The bipolar electrode is bent. (4) The bent bipolar electrode is attached to a fiber optic cannula. (**B**)A photograph of an assembled implant.

### Animals

All animal experiments were performed in accordance with the United Kingdom Animals (Scientific Procedures) Act of 1986 Home Office regulations and approved by the Home Office (PPL 70/8883). Three ChAT-IRES-Cre (JAX006410) mice were used (female, 8-37 weeks old) and housed individually in high-roofed cages with a 12 h:12 h light/dark cycle (light on hours: 7:00-19:00). Mice had *ad libitum* access to food and water. All experiments were performed during the light period. No blind and randomized experimental design was adopted due to the nature of the technical development study.

### Surgery

The surgical procedures have been described previously (Tsunematsu et al., 2019). Briefly, mice were anesthetized with isoflurane (5% for induction, 1-2% for maintenance) and placed in a stereotaxic apparatus (SR-5M-HT, Narishige). Body temperature was maintained at 37°C with a feedback temperature controller (40–90–8C, FHC). Lidocaine (2%, 0.1-0.3 mg) was administered subcutaneously at the site of incision. Two bone screws were implanted on the skull for monitoring cortical EEGs and twisted wires were inserted into the neck muscle for obtaining EMG signals. An additional bone screw was implanted over the cerebellum to provide a ground/reference channel. These electrodes were connected to a 2-by-3 piece connector (SLD-112-T-12, Samtec). Two additional anchor screws were implanted bilaterally over the parietal bone to provide stability and a small portion of a drinking straw was placed horizontally between the anchor screws and the connector. The viral vector (AAV5-CAG-flex-GCaMP6s-WPRE-SV40, Penn Vector Core; titer 8.3×10^12^ GC/ml) was microinjected (500 nl at 30 ml/min) (Nanoliter2010, WPI) to target the pedunculopontine tegmental nucleus (PPT) and laterodorsal tegmental nucleus (LDT) (-4.5 mm posterior, 1 mm lateral from bregma and 3.25 mm depth from brain surface). The micropipette was left in the brain for an additional 10 minutes and then slowly raised up. A hybrid implant (see above) was then implanted 3 mm deep from the surface of the brain and all components were secured to each other and the skull with dental cement.

### Recording procedures

After a recovery period (3-4 weeks), mice were habituated to being handled and tethered to the freely behaving system over several consecutive days. Mice were scruffed and the straw on the headcap slotted into a custom-made clamp, to keep the head still and absorb any vertical forces when connecting the electrophysiology and fiber photometry tethers to the headcap. Once connected, mice were placed in an open top Perspex box (21.5 cm × 47 cm × 20 cm depth) lined with absorbent paper, bedding and some baby food. During the habituation period, short recordings (20 – 30 minutes) were taken to test illumination parameters for the best signal to noise ratio. The illumination power was adjusted at the tip of optical fiber to 0.4 - 0.94 mW/mm^2^ for the 405 nm LED and 0.7 - 1.37 mW/mm^2^ for the 470 nm LED. Following the habituation period, simultaneous electrophysiological recording and calcium imaging was performed for 4-5 hours to allow for multiple sleep/wake transitions.

### Signal processing

All signal processing was performed offline using MATLAB (version 2018b, Mathworks).

#### Fiber photometry

Custom-written MATLAB scripts were used to compute fluorescent signals. To extract 405 and 470 nm signals, illumination periods were determined by detecting synchronization pulses. The median fluorescent signal was calculated during each illumination epoch (**Fig. 3B**). Because each illumination epoch consisted of pulses at 40 Hz, the fluorescent signals originally sampled at 1 kHz were effectively down-sampled to 40 Hz. Photobleaching was estimated by fitting a single exponential curve and the difference between the fluorescent signal trace and the estimate was further low-pass filtered at 4 Hz (**Fig. 3C**). To estimate moving artifacts, the filtered 405 nm signals were normalized based on the filtered 470 nm signals using a linear regression. To estimate fluorescent signals, the fitted 470 nm signals were subtracted from the scaled 405 nm signals (**Fig. 3D**).

**Figure 3.**
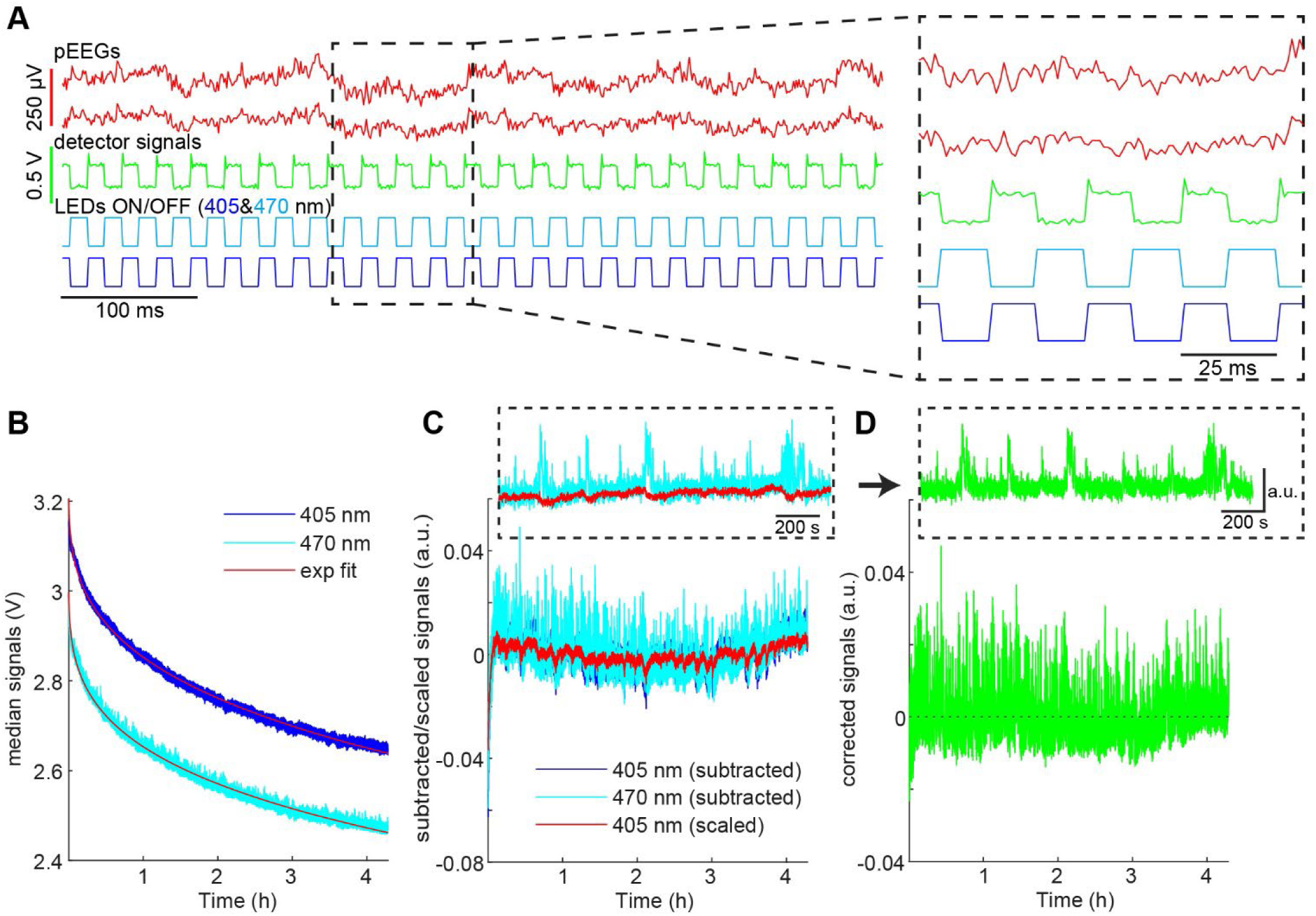
Signal processing. **(A)** Raw pontine EEGs (pEEGs) traces, photodetector signals, and LED illumination patterns. Note that no optical artifact was observed in pEEGs channels. **(B)** Median fluorescent signal values during each LED illumination step over a 4-hour recording period. *blue*, signals during 405 nm illumination. *light blue*, 470 nm illumination. *red*, exponential fit. **(C)** The median values in (**B**) were subtracted from the fitting curve (*blue*, 405 nm; *light blue*, 470 nm). The subtracted signals at 405 nm were linearly scaled (*red*) as artifacts. *inset*, an example fraction of 470 nm signals and scaled 405 nm signals. **(D)** The subtracted signals at 470 nm in (**C**) were corrected by subtracting signals from the scaled signals to provide normalized fluorescent signals. *inset*, an example fraction of the corrected signals.

#### Electrophysiology

Vigilance states were visually scored offline as described elsewhere (Tsunematsu et al., 2019). Wakefulness, NREM sleep, or REM sleep was determined over a 4-second resolution, based on cortical EEG and EMG signals using a custom-made MATLAB Graphical User Interface. The same individual scored all recordings for consistency.

To detect pontine waves (P-waves), two EEG signals in the pons were subtracted and filtered (5-30 Hz band-pass filter). If the signals cross a threshold, the event was recognized as pontine waves. To determine the detection threshold, a 1-minute segment of the subtracted signals was extracted from the longest NREM sleep episode to estimate stable noise level. The noise level was estimated by computing root-mean-square (RMS) values in every 10 ms time window. The threshold was defined as mean + 5 x the standard deviation of the RMS values. The timing of P-waves was defined as the timing of the negative peak. To generate surrogate P-wave timing during REM sleep (**Fig. 4C**), the number of P-waves during each REM sleep episode was held, but P-wave timing was randomly allocated during the episode. This surrogate timing was used to extract GCaMP6s signals for comparisons.

**Figure 4.**
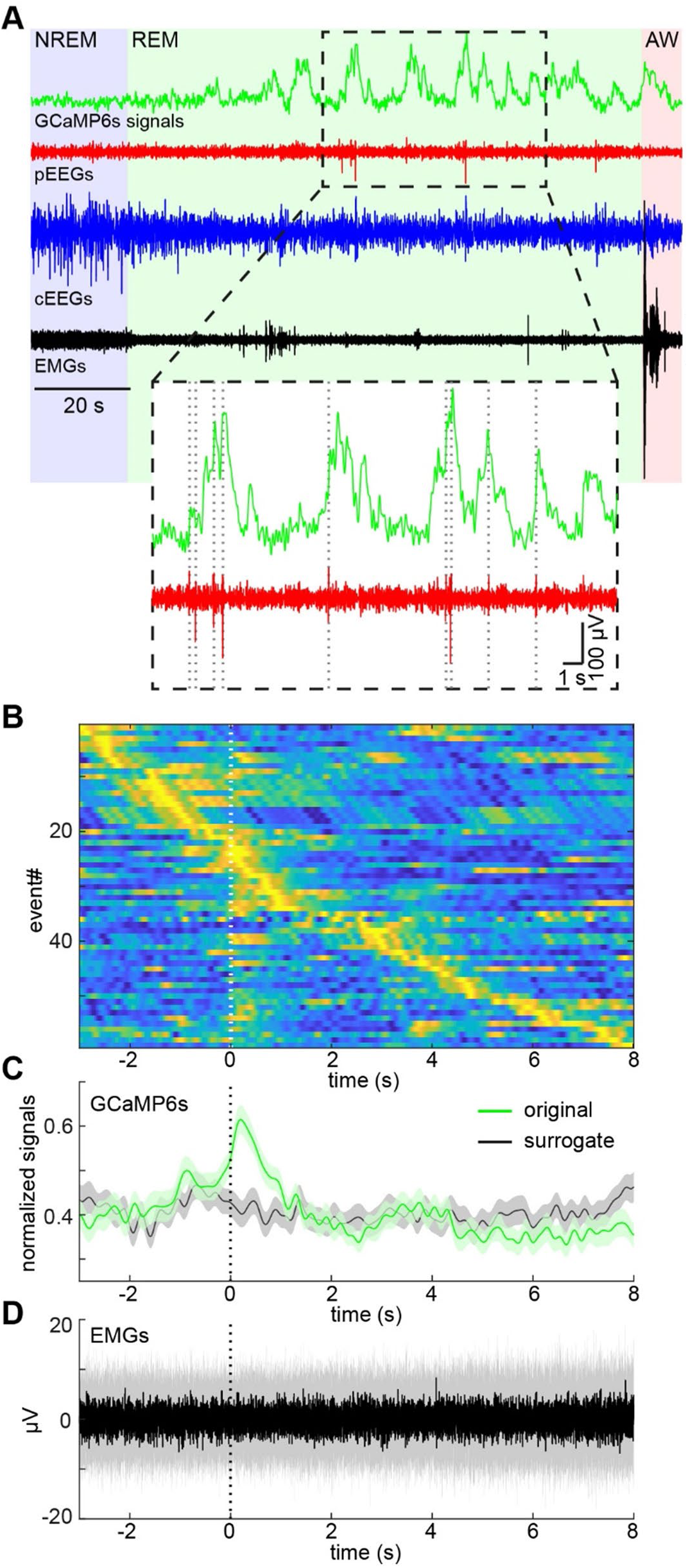
Pontine waves and cholinergic activity during REM sleep. **(A)** Example GCaMP6s, pontine EEGs (pEEGs), cortical EEGs (cEEGs) and EMGs signals around an example REM sleep episode. *Inset*, pontine waves (spiking events in pEEGs) often co-occurred with large activity of calcium signals. **(B)** P-wave-triggered calcium signals. GCaMP6s signals were aligned at the onset of pontine waves (P-waves) during REM sleep. GCaMP6s signals were normalized by the maximum value of each trace. **(C)** The mean profile of P-wave-triggered calcium signals. The normalized signals in (**B**) were averaged (green). In the surrogate condition (black), the timing of P-waves were randomly assigned during each REM sleep episode whilst still conserving the number of P-waves during the episode. Then the averaged calcium signals were computed. Errors, SEM. **(D)** The median profile of P-wave-triggered EMG signals. Shaded area, 25-75 percentile.

### Statistical analysis

Data was presented as mean ± SEM unless otherwise stated.

## Results

### Simultaneous monitoring of pontine EEGs and calcium transients in cholinergic neurons in freely behaving mice

We collected 6 datasets from 3 animals in the present study. The average recording duration was 253.5 ± 8.5 min (range, 212.7 – 267.1 min) (**Table 2**). **Figure 3A** shows representative raw traces of pontine EEGs, photodetector signals and LED illumination pulses. Our initial concern was that optical illumination might induce optical artifacts in pontine EEG signals as reported in optogenetic experiments (Kozai and Vazquez, 2015). However, no optical artifact was observed.

**Table 2.**
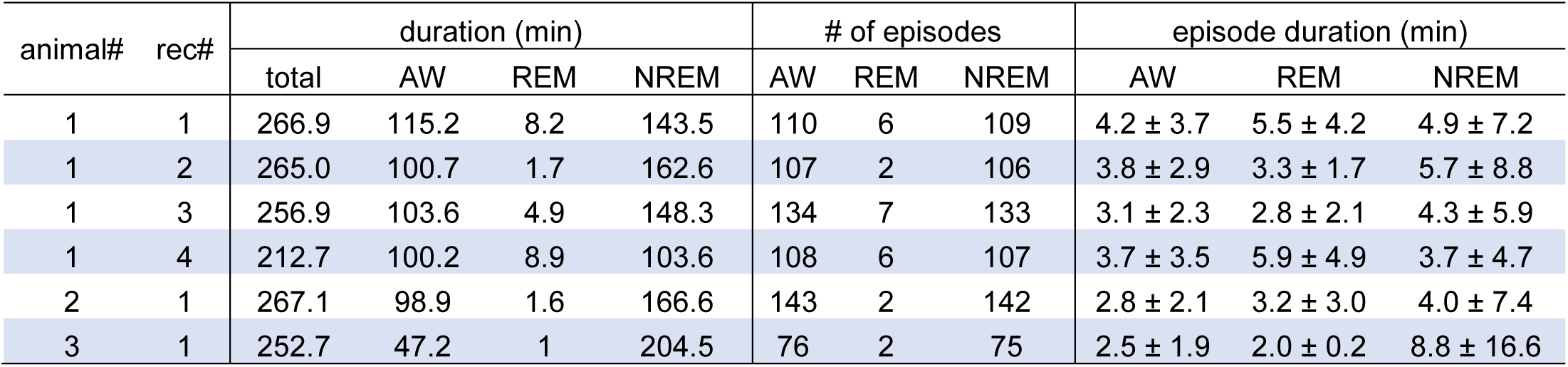
Statistics of sleep-wake cycles in individual recordings. AW, wakefulness; REM, rapid eye movement sleep; NREM, non-REM sleep. Data represents mean ± standard deviation.

To investigate neural activity across arousal states, it is crucial to monitor neural signals for several hours. Therefore, we also evaluated the stability of calcium transient amplitudes during the recording. While overall fluorescent signals decreased exponentially (**Fig. 3B**), calcium transients were relatively stable over several hours (**Fig. 3D**). Thus, our approach allows for the simultaneous monitoring both electrophysiological signals and calcium transients in freely behaving animals across the sleep-wake cycle.

### Pontine waves and calcium transients in pontine cholinergic neurons

In all six datasets, we observed multiple REM and non-REM (NREM) sleep episodes as summarized in **Table 2**. To demonstrate the potential of our approach, we tested the hypothesis that pontine (P) waves co-appear with calcium transients in pontine cholinergic neurons during REM sleep (Callaway et al., 1987;Datta, 1997). **Figure 4A** shows example signals around a REM sleep episode. We observed frequent calcium transients during REM sleep as well as phasic, large fluctuations of pEEGs. More importantly, these two events often co-occurred. To quantify this tendency, we detected P-waves and observed the associated calcium signals during REM sleep across recordings (**Figs. 4B and C**). Large calcium transients appeared around the timing of P-waves. To verify that these observations were not due to artifacts, we generated surrogate events where the timing of pseudo P-waves was randomly assigned during REM sleep whilst still maintaining the frequency of P-waves across REM sleep episodes. We then assessed the calcium transients around the surrogate P-wave events. As expected, the calcium transients would only be prominent around the onset of the real P-waves (**Fig. 4C**). In addition, we did not observe changes in EMGs associated with P-waves (**Fig. 4D**), indicating that these transients and P-waves were not due to movement artifacts. Thus, we confirmed that P-waves co-occurs with calcium transients in pontine cholinergic neurons during REM sleep.

## Discussion

A combination of electrophysiological recording and calcium imaging can be used to monitor cell-type-specific activity together with sub-second neuronal events in freely behaving animals. In this study, we utilized this approach to correlate calcium transients in pontine cholinergic neurons with P-waves during REM sleep in mice for the first time. The same approach can be applied in various experimental contexts. Thus, our method adds a novel tool to investigate state-dependent neural circuit dynamics *in vivo*.

Although there are a handful of commercially available fiber photometry systems, our system is easy-to-build and economical. All parts can be purchased from well-known suppliers and cost approximately 6,800 USD in total. Therefore, our system offers an affordable solution to integrate *in vivo* electrophysiology with calcium imaging in freely behaving rodents.

In the present study, we focused on the relationship between pontine cholinergic neurons and P-waves during REM sleep to demonstrate the capability of our system. Although P-waves or PGO-waves have been studied in several mammalian species, such as cats, monkey and rats, few studies have investigated P-waves in mice (Tsunematsu et al., 2019). Previous studies suggest that cholinergic neurons play a role in the induction of P-waves (Callaway et al., 1987;Steriade et al., 1990;Datta et al., 1992;Datta, 1997). In line with this, our results directly demonstrated that indeed cholinergic population activity co-occurs with P-waves for the first time. A limitation of GCaMPs is that it provides only an approximate reflection of neuronal spiking activity. Therefore, the exact temporal relationship between the firing of cholinergic neurons and P-waves still need to be investigated with the used of genetically encoded voltage indicators or electrophysiological techniques with optogenetic tagging. In addition to cholinergic neurons, it would be also interesting to monitor calcium transients in different cell types across pontine nuclei to characterize neural ensemble dynamics underlying P-waves.

A similar approach can be taken in different experimental settings. For example, field potentials can be monitored with cell type-specific calcium transients in task performing animals. Our system can be customized to add optogenetic stimulation by expressing red-shifted indicators and opsins sensitive to blue light (Chen et al., 2013). A bipolar electrode can be replaced by other types of electrodes to record broadband signals including spiking activity to correlate calcium transients with neuronal spiking. Fiber photometry system can be updated to utilize a tapered optic fiber to monitor activity in a larger area (Pisanello et al., 2019) or to perform cell-type-specific voltage imaging (Marshall et al., 2016;Kannan et al., 2018). In conclusion, our combinatory approach with electrophysiological recording and fiber photometry offers an affordable, but powerful solution to interrogate state-dependent neural circuit dynamics across various brain regions and behavioral states.

## Acknowledgements

This work was supported by BBSRC (BB/M00905X/1), Leverhulme Trust (RPG-2015-377), Alzheimer’s Research UK (ARUK-3033bb-CRT), and Action on Hearing Loss (S45) to SS.

## Authors contribution

AAP and SS designed and conceived the project. AAP and NM developed the recording system. AAP performed all experiments. AAP and SS analyzed data. AAP, NM and SS wrote the manuscript. KM and SS supervised NM and AAP, respectively.

